# A conserved dimerization motif squelches transcription

**DOI:** 10.64898/2026.11.16.767078

**Authors:** Julie M. Ray, Janet Solano Sanchez, Joseph O. Armstrong, Alexander R. Leydon, Jennifer L. Nemhauser

## Abstract

Squelching is an indirect inhibition of transcription that results from redistribution of cofactors away from DNA. Here, we describe a class of Lis homology (LisH)-derived peptides that act as transcriptional squelchers. LisH domains are a highly conserved motif found in 10s of thousands of proteins and are composed of ∼30 amino acids in a helix-turn-helix fold. By examining a phylogenetically diverse subset of LisH domains in a synthetic *Saccharomyces cerevisiae* assay, we identified a small number of hydrophobic residues within the first helix that are required for effective inhibition of reporter expression. In addition, squelching required binding to the general transcriptional cofactor Taf5, a subunit of the TFIID complex. Transcriptome-level impacts were largely correlated in yeast expressing two distinct LisH sequences, consistent with a model that LisH-based squelching results from a shared mechanism of sequestering Taf5. Consistent with that inference, overexpression of Taf5 partially reversed the squelching effect. Given the interest in squelching as a potential area for anti-cancer therapies, we also tested whether our shortest squelching sequence called YC5C-H1 had any effect on the human colorectal cancer cell line HCT-15. In addition to preferentially squelching high expression genes, YC5C-H1 sharply slowed growth of HCT-15 cells.

**Statement of Significance:** We have discovered that a class of peptides derived from the Lis homology (LisH) dimerization motif can inhibit transcription through association with the highly conserved general transcription cofactor Taf5, a subunit of the TFIID complex. The LisH peptides fit the established definition of a transcriptional squelcher, downregulating loci unable to compete for the limited cofactor in a dose-dependent manner.

## Introduction

Transcription of protein-coding genes requires coordination between regulatory transcription factors and the Pre-Initiation Complex (PIC) to load RNA polymerase II (Pol II) onto DNA (1). Most cofactors are present at limiting concentrations, and transcription factors provide specificity for where active transcriptional complexes assemble (2). Indirect repression, or “squelching”, can occur when cofactors are redistributed away from a regulatory region and are therefore unable to participate in transcription initiation (3). Squelching was first observed in *Saccharomyces cerevisiae* (yeast) when overexpression of the Gal4 activation domain (Gal4-AD) led to a decrease in transcription through sequestration of Mediator away from DNA (4).

Transplanting the auxin response pathway from Arabidopsis into yeast enabled testing of a range of hypotheses about transcriptional mechanisms, particularly in relation to corepressor function (5, 6, 7). In this synthetic system, a plant transcription factor activates a fluorescent reporter from a cognate regulatory region (8). As neither the activator nor its binding site has an exact counterpart in yeast, reporter output is insulated from endogenous regulation including compensation. We previously found that the first *α*-helix of LisH domains isolated from diverse eukaryotic proteins (hereafter referred to as H1s) were sufficient to repress the reporter in these assays (9). Reporter synthetic genetic arrays and proximity labeling connected the function of one of the LisH repression domains to direct physical interaction with core transcriptional machinery including Spt4 and Taf5 (5). From this result, we predicted that expressing the LisH alone, without the capacity to be recruited to DNA, might sequester cofactors away from promoters. Such an effect is the defining behavior of a transcriptional squelcher (4).

Here, we have adapted our synthetic yeast platform to test for squelching potential, and found several H1-containing peptides that could inhibit reporter expression. We focused our further analysis on one of these sequences, an H1 derived from the *Schizosaccharomyces pombe* protein YC5C (also known as Gid1). Using flow cytometry, transcriptomics, and protein-protein interaction assays, we found that transcription inhibition by YC5C-H1 in *S. cerevisiae* was dose-dependent and affected a specific subset of genes across the yeast transcriptome. The genomic pattern of inhibition was largely correlated with the effects of expressing a squelching peptide from Arabidopsis, pointing to a likely shared mechanism of action. Recently, the therapeutic potential of squelching was demonstrated when a peptide targeted to the histone fold domain of TFIID proteins Taf4 and Taf12 significantly slowed growth of myeloid leukemia cells (10). To test the effects of YC5C-H1 on cells with aberrant transcription, we tested for squelching effects on the human colorectal cancer cell line HCT-15. In addition to slowed growth, transcriptomic analysis of HCT-15 cells expressing YC5C-H1 revealed significant downregulation of cell cycle regulators and genes associated with protein folding.

## Results

### A conserved dimerization domain can squelch transcription when not recruited to DNA

In the recapitulated auxin response circuit in yeast, an ARF transcription factor binds to a promoter upstream of a fluorescent reporter (ARF-reporter). In the absence of auxin, ARF activity is repressed via interaction with an adaptor protein that can be fused to a corepressor (7) or expressed independently (5). Using this circuit as a synthetic repression assay, we identified an autonomous repression domain in the first helix of the LisH domain in the plant corepressor TPL, which we refer to as H1. Functional characterization of H1s across extensive sequence and functional context revealed that many H1s retained repression activity (9).

To adapt this assay to test squelching, we expressed candidate peptides in the context of the simpler ARF-reporter activation module where no corepressor or adaptor protein are present. We defined squelching as decreased reporter fluorescence that was quantified by flow cytometry. Three H1s with clear repression activity (TPL-H1, YC5C-H1, and LUG-H1) (9) significantly squelched the reporter relative to the control (Figure 1a). GID8-H1, a quite weak repressor despite relatively high protein accumulation (9), was also a weak squelcher. YC5C-H1 accumulated to the highest level of the strong squelchers we tested (Figure 1a). As seen previously for Gal4-AD (4), there was a direct correlation between YC5C-H1 dosage and squelching strength (Figure 1b).

**Fig. 1.**
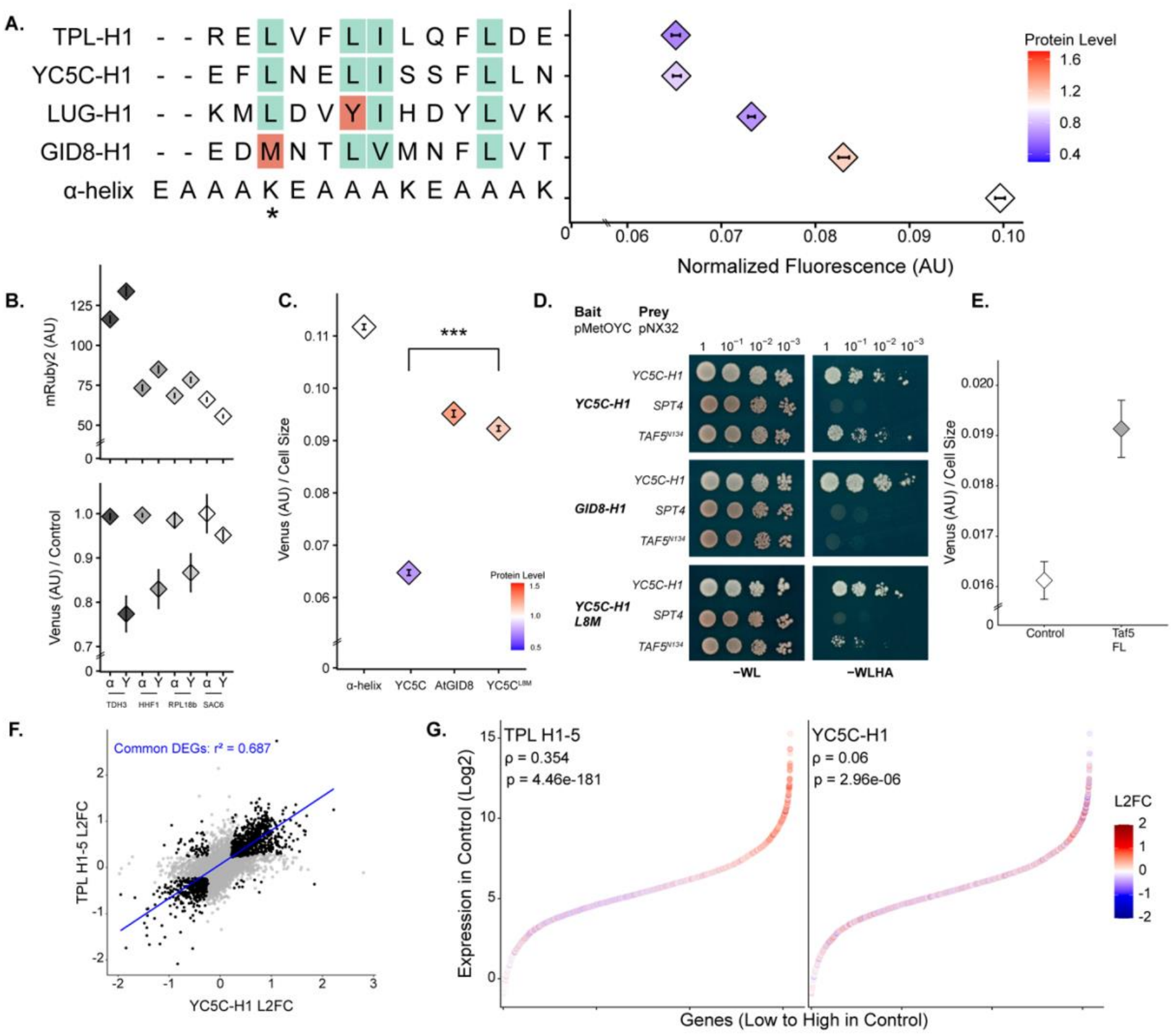
LisH peptides squelch transcription in a sequence- and dose-dependent manner through Taf5 and reshape the yeast transcriptome. All experiments were performed in *Saccharomyces cerevisiae*. Throughout, Venus fluorescence reports transcriptional output from the synthetic ARF19-responsive promoter and mRuby2 fluorescence reports H1 protein accumulation, both in arbitrary units (AU). (A) Flow cytometry quantifying squelching by LisH H1s. Venus fluorescence is normalized to cell size (FSC-A) and diamonds are colored by protein accumulation, measured as mRuby2 fluorescence normalized to the α-helix control. Conserved hydrophobic residues in the aligned H1 sequences are shaded cyan and natural variation at these positions red; the asterisk marks the position substituted in YC5C-H1^L8M^. (B) mRuby2 accumulation plotted above Venus fluorescence normalized to the α-helix control, for YC5C-H1 expressed from promoters of decreasing strength (pTDH3, pHHF1, pRPL18b and pSAC6, strongest to weakest). α, α-helix control; Y, YC5C-H1. Squelching strength scales with YC5C-H1 accumulation. (C) Venus fluorescence normalized to cell size for the α-helix control, YC5C-H1, GID8-H1 and YC5C-H1^L8M^, colored by protein accumulation as in (A). The weak squelcher GID8-H1 and the YC5C-H1^L8M^ substitution are both impaired in squelching. ***P < 0.001. (D) CytoSUS assays testing interaction between each H1 bait (pMetOYC) and three preys (pNX32): YC5C-H1, SPT4 and Taf5^N134^, the N-terminal 134 amino acids of ScTaf5 containing the LisH domain. Cultures were spotted in tenfold dilutions from undiluted to 10^<sup>−</sup>3^. – WL indicates growth media lacking tryptophan and leucine, which selects for yeast carrying both bait and prey constructs without testing for interaction between them; –WLHA indicates media lacking tryptophan, leucine, histidine and adenine, in which only yeast expressing interacting bait and prey can grow. All three baits interact with the YC5C-H1 prey, consistent with LisH-mediated dimerization, and none interacts with SPT4. YC5C-H1 interacts with Taf5^N134^, while GID8-H1 and YC5C-H1^L8M^ show no or much reduced interaction with Taf5. (E) Complementation of YC5C-H1 squelching by full-length ScTaf5. Venus fluorescence, normalized to cell size (FSC-A), in strains co-expressing YC5C-H1 with full-length Taf5 (Taf5 FL) or with an empty NubG prey (Control); full-length Taf5 rescues reporter expression. Each replicate is the median Venus/FSC-A of ∼10,000 gated events; diamonds, mean of three biological replicates; bars, SD. (F) Scatterplot of all genes, with significant DEGs shared between squelchers in black and the remainder in gray. YC5C-H1 log2 fold change on the x-axis, TPL H1-5 log2 fold change on the y-axis. Blue line, line of best fit for significant DEGs shared between squelchers: y = 0.068 + 0.74x, R^2^ = 0.687. (G) Significant DEGs plotted by expression in the control strain (x-axis, genes ordered by control expression; y-axis, log2 of control counts per million) and colored by log2 fold change (L2FC). Left, TPL H1-5; right, YC5C-H1. ρ and P, Spearman correlation between fold change and expression in the control strain. Scale: L2FC –2 to +2, identical for both.

TPL-H1 and YC5C-H1 were the strongest squelchers and shared four conserved hydrophobic residues, predicted to align along a common face of the helix (Figure 1a), which were less well conserved in the weaker squelchers. The weakest squelcher GID8-H1 was missing one of these highly conserved leucine residues, and a YC5C-H1^L8M^ mutant was severely impaired in its ability to squelch (Figure 1c). Mutating the equivalent face of the Taf5 LisH helix also abolishes TPL-mediated repression (5), consistent with squelching requiring an intact LisH–LisH interface. Consistent with the mechanism previously established for TPL (11), cytoplasmic split-ubiquitin (cytoSUS) assays revealed that YC5C-H1 interacted with an N-terminal truncation of yeast Taf5 containing a LisH domain (Taf5N134) (Figure 1d). The weak squelchers GID8-H1 and YC5C-H1^L8M^ showed no or much reduced interaction with Taf5N134 (Figure 1d), consistent with a model that squelching acts by sequestering Taf5. To directly test this hypothesis, we co-expressed YC5C-H1 with full-length ScTaf5, and found that increasing the abundance of the cofactor was able to counteract the effects of squelching (Figure 1e).

If squelching by H1s is acting through sequestering Taf5, we predicted that we should see similar patterns of impacted genes genome-wide regardless of which H1 was expressed. Moreover, squelching is not predicted to affect all genes equally; rather, it should impact the subset of genes that are most sensitive to the reduction in availability of a transcriptional cofactor (3). To test these predictions, we analyzed the transcriptome of strains expressing either TPL H1-5 or YC5C-H1 (SI Appendix, Fig. S1; Dataset S1). Our results were consistent with transcription inhibition by squelching. First, differentially expressed genes (DEGs) correlated in both sign and magnitude between TPL H1-5 and YC5C-H1 (Figure 1f, blue trend line, R^2^ of 0.687; Dataset S1). Second, we found that genes expressed at low levels in our control condition were the most affected by expression on TPL H1-5 (Figure 1g, left), and a similar trend could be seen for YC5C-H1 with the addition of a cluster genes in the highest control expression quartile that were not affected by TPL H1-5 (Figure 1g, right; Dataset S1).

### YC5C-H1 slows proliferation and downregulates cell cycle and protein folding programs in human cancer cells

Squelching has previously been shown to slow the growth of cancer cells (10), and we wondered whether expression of YC5C-H1 would act similarly. When YC5C-H1 was stably expressed in the human colorectal cancer line HCT-15, the cells showed striking changes to morphology (Figure 2a-d) and significantly inhibited growth (Figure 2e). Cell size was also impacted (Figure 2f, SI Appendix, Fig. S2b, right). Staining DNA of control and squelched cells revealed a significant enrichment of squelched cells in G2 (SI Appendix, Fig. S2b, left), with no significant change in other phases.

**Fig. 2.**
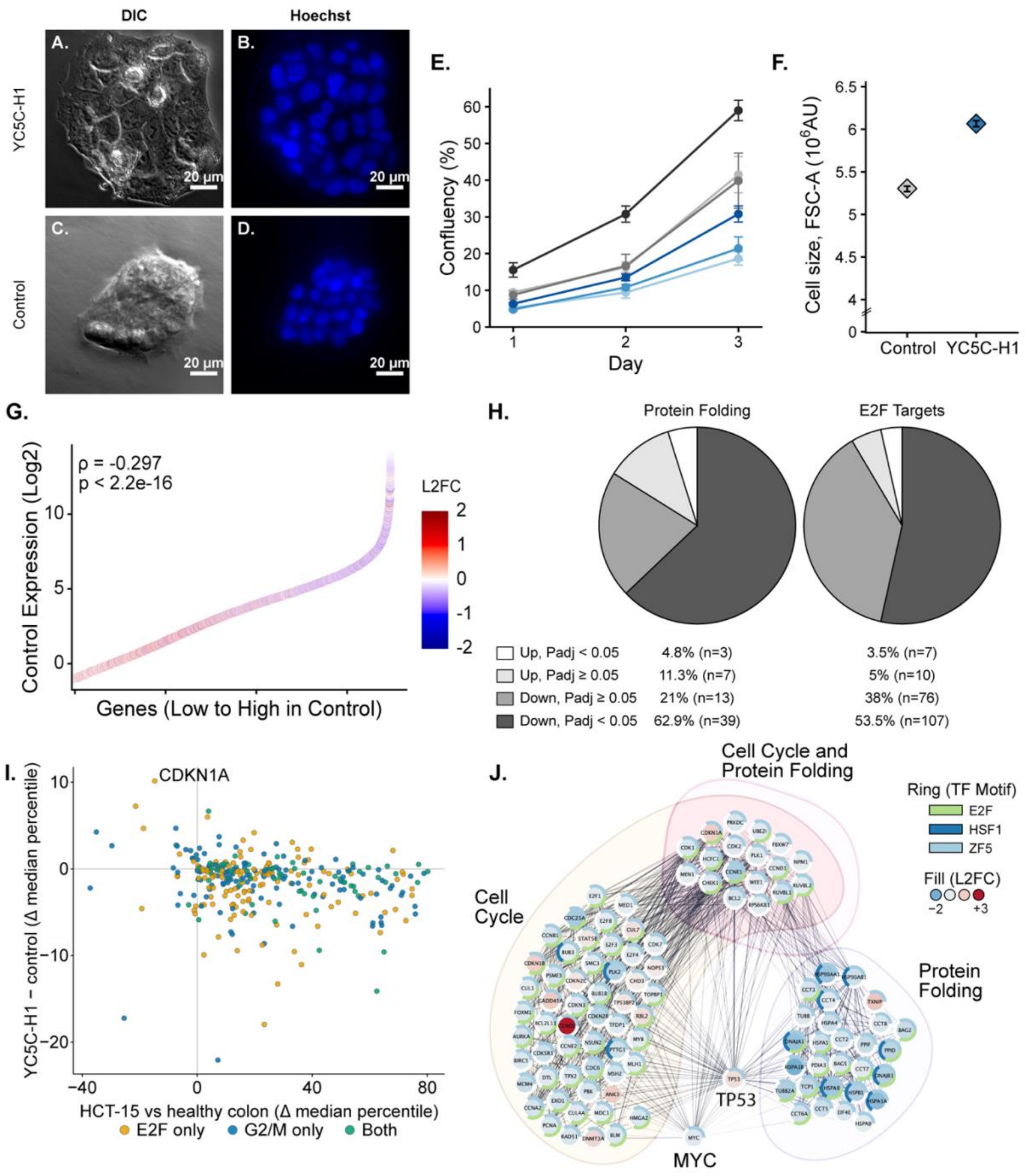
YC5C-H1 slows proliferation and downregulates cell cycle and protein folding programs in HCT-15 cells. All panels show the human colorectal cancer line HCT-15, comparing the control line with the YC5C-H1-expressing (squelched) line. RNA sequencing was performed on three biological replicates per cell line. (A) DIC microscopy of squelched HCT-15 cells. Scale bar, 20 µm. (B) Hoechst stain microscopy of squelched HCT-15 cells. Scale bar, 20 µm. (C) DIC microscopy of control HCT-15 cells. Scale bar, 20 µm. (D) Hoechst stain microscopy of control HCT-15 cells. Scale bar, 20 µm. (E) Confluency time course for control (gray) and squelched (blue) HCT-15 cells, measured from day 1. Line darkness corresponds to seeding density, from 10,000 cells per well (faintest) to 30,000 cells per well (darkest). Average confluency from three biological replicates per cell line at each seeding density are shown. Error bars are the standard error of the mean across three replicate wells, each the average of three imaging fields. (F) Cell size (FSC-A, AU) of control versus squelched HCT-15 cells measured by flow cytometry; three biological replicates per cell line. Diamonds, median of pooled events; error bars, bootstrapped 95% CI. (G) Spearman’s correlation of differential gene expression in the squelched line against expression level in control HCT-15 cells. Rho = –0.297, P < 2.2 × 10<sup>−</sup>^16^. (H) Left: differential expression of protein folding chaperone genes, grouped as up with Padj < 0.05, up with Padj ≥ 0.05, down with Padj ≥ 0.05 and down with Padj < 0.05. Right: differential expression of E2F target genes, grouped in the same categories. (I) Scatterplot of G2/M-associated genes (blue) and E2F target genes (yellow), with genes in both sets in green. X-axis, expression in control HCT-15 cells relative to healthy colon tissue (Δ median percentile); y-axis, differential expression in the squelched versus control line (Δ median percentile). (J) STRING network of cell cycle and protein folding genes. Nodes are filled by log2 fold change (range –2 to +3) and outlined with a split donut indicating predicted regulation by ZF5 (light blue), HSF1 (dark blue) and E2F (light green). Shaded envelopes group nodes by shared STRING edges, which represent predicted functional interactions: cell cycle only, cell cycle and protein folding, and protein folding only, as labeled.

Transcriptome analysis of these cells revealed that YC5C-H1 expression had the largest impact on genes in the top quartile of expression in HCT-15 cells (Figure 2g), a contrast to what we observed in yeast. GO analysis on highly significant squelched DEGs (padj < 0.001) identified an enrichment in GO terms associated with protein folding as well as several terms associated with the cell cycle, including mitosis and cell cycle phase transition (Dataset S2). In the downregulated DEGs, there were a significant enrichment of cyclins and CDK activators, as well as terms associated with the TP53 signaling pathway (Dataset S2; SI Appendix, Fig. S2a). We observed squelching of many highly expressed protein folding chaperones, including several HSPs and components of the TriC/CCT complex (SI Appendix, Fig. S2c; Dataset S2). Analysis of TF binding motifs in squelched genes identified an enrichment of E2F target genes, a transcription factor known to regulate cell cycle phase transition (Dataset S2). GO analysis on DEGs across all genes annotated to protein folding and to regulation by E2F found that approximately 64% of protein folding genes were significantly downregulated (84% down without filtering for significance), and 53% of E2F target genes were significantly downregulated (91% down without filtering) (Figure 2h).

We next extracted a list of G2/M and E2F cancer hallmark genes and found, as expected, that these genes were largely upregulated in HCT-15 cells compared to healthy colon tissue (SI Appendix, Fig. S2d). In HCT-15 cells expressing YC5C-H1, the majority of these genes were downregulated (Figure 2i), including a subset returned to the expression range of healthy colon tissue (SI Appendix, Fig. S2e). Using Gene Set Enrichment Analysis against the Genotype-Tissue Expression database, we found the G2/M transition to be among the most significantly affected programs by YC5C-H1 expression (NES = −2.89, padj = 1.2 × 10<sup>−</sup>^27^), with associated programs downstream of E2F and MYC also significantly downregulated (Dataset S2). A STRING network analysis on highly significant genes associated with cell cycle and protein folding identified TP53 and MYC as hubs, with several high-confidence edges connecting HSP and TCP/CCT genes to cell cycle genes through them (Figure 2j). TP53 was significantly upregulated while MYC was significantly downregulated. We identified an enrichment of binding sites for ZF5, a ubiquitous zinc finger transcription factor predicted to bind GC-rich promoters (12), as well as for E2F and HSF1. Nearly all genes in the network contained a ZF5 motif and many in all three groups contained an E2F motif, while a subset of protein folding genes and a smaller subset of cell cycle genes contained a well-annotated HSF1 motif. Nearly all genes in the network were downregulated with the exception of one cyclin and a cell cycle inhibitor.

## Discussion

Many LisH domains can repress transcription (9), and this activity has been shown to require nucleating components of the transcription initiation complex for a LisH-containing corepressor from plants (5, 6). Leveraging a synthetic, orthogonal transcriptional assay in yeast, we tested whether LisH-derived peptides could lead to similar transcriptional interference through sequestration of transcriptional cofactors as originally identified in the squelcher Gal4-AD (4). We found that LisH-containing peptides could inhibit transcription, and that this squelching effect required conservation of residues in the predicted hydrophobic surface of H1 and interaction with the general cofactor Taf5. Expression of two different LisH-based sequences impacted a largely overlapping group of yeast genes, likely those less able to compete for the more limited pool of TFIID complexes. Consistent with the impact seen by other proteins that sequester TFIID cofactors, YC5C-H1 inhibited growth of human cancer cells and downregulated a number of cell cycle-associated genes.

LisH domains have now been shown to block transcription in two distinct ways. In the context of a full-length corepressor, they can block transcription initiation when recruited to a specific locus. This mode of repression has been termed “primed”, as it appears to lead to assembly of an inactive pre-initiation complex that can rapidly transition to active transcription once a signal is received (13). In the second mode described here, a LisH-containing peptide that is unable to be recruited to DNA binds to the same cofactors as the corepressor but sequesters them away from the genome. This second mode is consistent with the established definition of squelching.

Rather than a generic repression of all transcription (3), our results suggest that the squelching effect is specific to reducing availability of Taf5, and that it therefore impacts the loci least able to compete for a limited supply of TFIID complexes. Consistent with this interpretation, we were able to pinpoint residues required for the squelching effect (Figure 1c), and found that these were the same residues required for binding to Taf5 (Figure 1d). Squelching also scaled with YC5C-H1 expression level, consistent with a model of competition for a limited pool of specific interaction partners (Figure 1b).

Beyond their utility in probing fundamental features of transcriptional regulation, peptide squelchers are also potentially useful additions to the synthetic biology toolbox. Connecting synthetic circuit performance with repression of squelcher expression (or induction of a squelcher depending on the selection scheme) could serve as a sensitive, dose-dependent output. The dose responsiveness of squelching falls within a reasonable range of promoter strengths, and, depending on the experimental design, may provide improved dynamic range when compared with currently favored outputs like prototrophy, antibiotic resistance or fluorescence. For example, such an output could be quite useful in Deep Mutational Scanning in microbes or mammalian cells (14, 15). It could also be useful in iterative cycles of optimization of synthetic circuits (16).

The effect of squelching on HCT-15 cells suggests other potential applications. The morphological changes we observed in squelched HCT-15 cells, namely enlargement, altered shape and an apparent increase in nuclear volume, together with accumulation of cells in G2 and upregulation of TP53 and CDKN1A, resemble a senescence-like state (17). A similar phenotype has been described in HCT-116 colon cancer cells treated with the bacterial cyclomodulin CNF1 (18). CNF1-induced senescence leads to cells accumulating at G2/M, increases rates of endoreplication and cell enlargement, and leads to upregulation of p53, p21 and p16. Similarly, E2F1 is known to suppress senescence in colorectal cancer cells (19). The broad downregulation of E2F target genes we observe (Figure 2h) would be expected to relieve this suppression—an exciting interpretation to test in future studies.

## Materials and Methods

### Cloning

Vectors used for yeast flow cytometry experiments were generated with the Yeast Molecular Cloning Toolkit (20), using a combination of site-directed mutagenesis and Golden Gate assembly. LisH H1 constructs were derived from the *A. thaliana* genes *TPL* (AT1G15750), *LUG* (AT4G32551), and *GID8* (AT1G61150), while *YC5C-H1* was cloned from the *S. pombe* gene *GID1* (SPCC1259.12c). The aligned H1 sequences are shown in Figure 1a and were described in prior work (9). H1s were constructed using the leader sequence “MSSLS” immediately followed by the H1 sequence, a small flexible linker (GRGPGGGHQ), and a 1xHA tag. This construction was used for strains used in RNA-seq. To overcome issues with protein stability among H1s with sequence diversity, an mRuby2 tag was added after the HA tag using Golden Gate. Flow cytometry to measure mRuby2 fluorescence was used to quantify protein in H1-mRuby2 strains. Bait vectors used for cytoSUS were constructed using site-directed mutagenesis to generate pMetOYC driving YC5C-H1, GID8-H1, and YC5C-H1^L8M^. The prey vector pNX32:YC5C-H1 was constructed using the same method.

### Yeast methods

#### Strain preparation and growth

Standard synthetic dropout (SDO) and yeast extract-peptone-dextrose plus adenine (YPAD) media were used to grow strains, with care taken to use the same batch for related experiments. Yeast transformations were performed using a standard LiOAc transformation protocol. All strains were kept under constant dropout selection on SDO (-Uracil) in a background containing an integrated Single Plasmid Auxin Response Circuit (SPARC). Construction of the SPARC was described previously (6). CytoSUS bait and prey constructs were transformed into MATa (selected on SDO –L) and MATα (selected on SDO –W) yeast strains respectively, then mated overnight in 2mL YPAD media at 30°C with 100rpm shaking. Mated cultures were plated the following day on SDO –WL to select for diploids. Strains were re-struck after mating before use in cytoSUS experiments. Complete Supplement Mixture (CSM) media was used subsequently to grow liquid cultures and plate cytoSUS strains.

#### Flow cytometry sample preparation and data collection

Flow cytometry strains were grown in 1mL SDO –U media overnight at 30°C with 350rpm shaking in deep 96-well plates covered with breathable tape. Cultures were grown overnight until confluent, diluted back, and grown 4 hours to capture all strains in the logarithmic growth phase. After 4 hours, mVenus fluorescence and cell size were measured using a Becton Dickinson (BD) special order cytometer with a 514 nm laser exciting fluorescence that is cut off at 525 nm prior to photomultiplier tube collection (BD). mRuby2 fluorescence was measured using a SONY 3800 Cytometer (excitation 561nm, emission 605nm).

#### Flow cytometry data analysis

Events were annotated and subset to singlet yeast using the FlowTime R package. A total of 10,000–20,000 events above a 400,000 FSC-H threshold (to exclude debris) were collected for each sample and data were exported as FCS 3.0 files for processing using the flowCore R software package and custom R scripts (8, 21). Analysis and visualization were performed in R, and all flow cytometry experiments are plotted with 95% confidence intervals. For all flow experiments, data from at least two independent replicates were combined and plotted in R using ggplot2. Flow cytometry experiments for LisH H1s (Figure 1a) and the YC5C-H1^L8M^ mutant (Figure 1c) were performed in biological quadruplicate, YC5C-H1 dosage experiments (Figure 1b), and flow cytometry on strains used for RNA-sequencing were performed in biological duplicate.

#### RNA extraction and sequencing

Cultures were grown in biological duplicate in a volume of 5mL SDO –U overnight at 30°C with 250rpm shaking. Cultures were diluted back the following morning, and flow cytometry was performed on mid-log cultures after 4 hours as described above, using only the BD Accuri C6 cytometer. ∼100 million cells per culture were immediately harvested following flow cytometry and RNA was extracted using the Yeastar RNA kit. RNA was shipped to Azenta (GENEWIZ) for quality control, renormalization, addition of an ERCC spike-in control, and bulk RNA sequencing.

#### CytoSUS protein interaction assay

We performed protein-protein interaction assays using a modified version of the cytoplasmic split-ubiquitin system (11) which we have described previously (6). Diploid strains were grown overnight in 2mL CSM –WL +250µM Methionine at 30°C with 220rpm shaking. Culture density was measured the following morning by measuring the OD_600_ of a 1:50 culture dilution. Cultures were normalized to 1OD in 200µL of sterile dH_2_O and serially diluted in a 96-well plate. Strains were plated on CSM –WL and CSM –WLHA (+250µM Methionine) plates and allowed to grow for 2 days in a 30°C incubator. Plates were imaged using an Epson V800 Perfection flat-bed scanner.

#### Taf5 complementation flow cytometry

Strains carrying pTDH3-YC5C-AFB2-ARF19 were mated to strains carrying either pNX-TAF5-FL or the empty NubG prey pNX32-NUB-3xHA as described above for cytoSUS, and diploids were selected on SDO lacking uracil, tryptophan and leucine. Cultures were grown and sampled in logarithmic phase as described above. Venus fluorescence was measured on a BD Accuri C6 flow cytometer (FL2). Three independent biological replicates, each started from a separate single colony, were collected per genotype.

## Statistical Analysis

### Yeast RNA-seq data analysis

Transcript counts per million (TPM) were processed with DESeq2 (22) to generate mRNA count tables, Log_2_ Fold Change of DEGs, and adjusted DEG significance values with cutoff p<0.05 (23). Spearman’s correlations were performed on all transcripts with a TPM score (transcripts per million) over 0.5. We indexed the list of significant DEGs based on their expression in WT (the mRuby2 control) and performed a Spearman’s correlation analysis across all DEGs in R. Spearman’s analysis results were superimposed onto a reverse logarithmic curve (log_2_(mean mRuby2 counts)) of DEGs colored by their log_2_ fold change in the squelching condition. All transcripts were mapped in the correlation analysis, then colored based on their significance in either condition (padj < 0.05).

### *HCT-15 RNA-seq data* analysis

RNA sequencing data from HCT-15 cells was performed using an identical DEseq2 pipeline to yeast. Spearman’s correlation analysis was performed both across the curve (unfiltered DEGs with a TPM > 0.5) and in quartiles, computing rho and p-values for each quartile of gene expression. GO analyses were performed in gProfiler on downregulated DEGs with a strict significance cutoff (padj < 0.001), selecting for GO: Biological Process terms. GO terms were manually curated to highlight the most enriched terms, as well as excluding generic or repetitive terms. GO analysis was performed in parallel using the NIH DAVID Bioinformatics database (24), performing functional analysis and clustering. Differential expression pie charts for protein folding genes and E2F targets were generated from L2FC data on all genes within the GO term and separated by the significance of their L2FC in the squelching condition.

### Tissue expression database analyses

Oncogene panel analysis was performed using the COSMIC Cancer Gene Census, selecting for hallmark colorectal cancer genes (25); genes were classified as oncogenes or tumor suppressors based on curated CRC role annotations. For each gene, baseline expression in healthy colon tissue was drawn from the GTEx database.

Hallmark CRC genes were extracted from DEGs (without significance thresholds, to map the maximum number of genes in our dataset). For each gene, we computed the difference in median within-sample expression percentile between HCT-15 cells and healthy colon (GTEx, 26) on the x-axis, and between YC5C-treated and control HCT-15 cells on the y-axis; within-sample percentile ranking was used so that the HCT-15 RNA-seq and GTEx data could be compared despite different normalization. The same analysis was performed using the Hallmark cell cycle (G2M checkpoint) and E2F target gene sets from Molecular Signatures Database (MSigDB) (27, 28). Pre-ranked gene set enrichment analysis (GSEA) was performed to test whether proliferation programs were coordinated by YC5C-H1. Genes were ranked by the DESeq2 Wald statistic for YC5C-H1 versus control, and enrichment against the MSigDB Hallmark collection was computed with the fgsea R package (minSize 10, maxSize 500). Normalized enrichment scores (NES) and Benjamini–Hochberg-adjusted p-values were reported across all 50 Hallmark sets.

### STRING Network

The STRING network (29) was constructed in Cytoscape (30) using differentially expressed genes (padj < 0.001). Cell cycle and protein folding genes were selected and clustered based on functional enrichment for corresponding GO Biological Process terms. Genes were clustered with a permissive granularity cutoff of 1 and another functional enrichment was performed on the clustered subset, searching for highly significant Transfac Transcription Factor binding motifs. Sub-groups and hubs were manually separated based on shared edges and color coded by their group.

Claude 5 Sonnet was used for troubleshooting and debugging R code.

### Human Cell Methods

#### Cell line and culture

HCT-15 human colorectal adenocarcinoma cells (ATCC, CCL-225) were maintained in RPMI-1640 supplemented with 10% fetal bovine serum and 1× antibiotic-antimycotic (ABAM) at 37 °C in a humidified incubator with 5% CO<sub>2</sub>and were passaged using TrypLE.

#### Expression constructs and lentiviral transduction

The YC5C-H1:eGFP expression construct and the eGFP-only control construct were synthesized by Epoch Life Sciences and propagated in NEB Stable cells at 30 °C; both were carried on a lentiviral transfer plasmid bearing a puromycin-resistance cassette. Lentiviral particles were produced in LV-MAX Production Cells (a HEK293-derived suspension line; Thermo Fisher) by co-transfection of the transfer plasmid with the second-generation packaging and envelope plasmids psPAX2 and pMD2.G using the LV-MAX Transfection Kit (Thermo Fisher), and 5 mL of viral supernatant was collected. HCT-15 cells were transduced by spinoculation in a 6-well plate. Each transduction well received a titration of viral supernatant in 3 mL total RPMI together with a 10 µg/mL polybrene; a medium-only well (no virus, no polybrene) and a polybrene-only well (no virus) were included as controls. The plate was partially sealed with tape, centrifuged at 1,000 × g for 2 h at 30 °C, and returned to the incubator; cells were allowed to recover for 4–24 h at 37 °C before the medium was replaced.

#### Selection

Beginning 2 days post-transduction, transduced cells were placed under puromycin selection. The puromycin concentration was increased stepwise over several days to a maximum of 3 µg/mL, and cells were maintained under selection until all non-transduced cells in a parallel control well had died.

#### Fluorescence-activated cell sorting

Following puromycin selection, eGFP-expressing cells were bulk-sorted by fluorescence-activated cell sorting on a Sony SH800 cell sorter (Sony Biotechnology, San Jose, CA), collecting eGFP-positive cells relative to non-transduced controls after gating on forward- and side-scatter to exclude debris and doublets. The resulting polyclonal populations were returned to culture and expanded for subsequent experiments.

#### Growth (confluency) time course

For the growth assay (Figure 2e), cells were seeded into 24-well plates at 10,000, 20,000, and 30,000 cells per well, with three wells per seeding density, and imaged every 24 h over 4 days on a Nikon Ti-E inverted microscope equipped with a Plan Fluor 10×/0.30 NA dry objective and an Andor iXon EMCCD camera (1.32 µm/pixel), controlled by Micro-Manager 2.0.3. Transmitted light and GFP (Ex480) channels were acquired at 50 ms and 200 ms exposure, respectively. Confluency was quantified as cell-free (“free”) area in ImageJ/Fiji. The growth assay was performed with three biological replicates per cell line at each seeding density.

#### Live-cell imaging

HCT-15 cells stably expressing YC5C-H1:eGFP or eGFP alone were seeded in glass-bottom plates and labeled with Hoechst 33342 to mark nuclei prior to imaging. Cells were imaged on an inverted Nikon Ti microscope with a Plan Apo 100×/1.45 NA oil-immersion objective and a cooled Andor iXon EMCCD camera (model DU8201_BV), controlled through Micro-Manager 2.0.3, and were maintained at 37 °C and 5% CO<sub>2</sub>in a stage-top incubator throughout imaging. For each field, nuclei were imaged by Hoechst 33342 fluorescence (365 nm excitation at 10% LED intensity from a Sutter Lambda 721 LED source, DAPI filter cube, 50 ms) alongside a transmitted-light DIC image (50 ms), both acquired as single focal planes at a pixel size of 0.13 µm. For each cell line, 10 fields were imaged across 3 wells in a single session. Images were processed and analyzed in Fiji/ImageJ. Significant differences were determined using analysis of variance (ANOVA) with a post-hoc Tukey’s honest significance difference test.

#### RNA extraction and sequencing

Cells were seeded at 3.5 × 105 cells per well in 6-well plates and grown for 2 days; three biological replicates per line were harvested for RNA extraction. Total RNA was extracted using the Monarch Total RNA Miniprep Kit (New England Biolabs), quantified on a NanoDrop spectrophotometer, and normalized to 50 ng/µL in 50 µL aliquots. RNA was shipped to Azenta (GENEWIZ) for quality control, renormalization, addition of an ERCC spike-in control, and bulk RNA sequencing.

#### Cell cycle analysis

Cell cycle distribution was assessed by propidium iodide (PI) staining. Cells were harvested at ∼50% confluency, resuspended in 1 mL cold PBS, and pelleted by centrifugation at 4 °C. The pellet was resuspended in 300 µL cold PBS, and cells were fixed by adding 3 mL of ice-cold 70% ethanol dropwise with gentle inversion, then incubated on ice for 1h. Fixed cells were rehydrated and washed in PBS, then resuspended in a PI/RNase staining buffer prepared by PI and RNase A to PBS (final concentrations ∼50 µg/mL PI and 100 µg/mL RNase A); RNase treatment ensured PI stained DNA only. Cells were incubated for 30 min at room temperature in the dark and analyzed by flow cytometry on a BD Accuri C6 Plus. PI intensity (FL2-A) was used to assign cell cycle phase, with 2n and 4n peaks corresponding to G1 and G2/M, respectively; cell size was computed from FSC-A of the same gated events. The assay was performed with three biological replicates per cell line.

## Data Availability Statement

Yeast and human RNA-seq data have been deposited in NCBI Gene Expression Omnibus under accession GSE354717 and will be publicly available as of 20 November 2026. Flow cytometry data, plasmid sequences, and analysis code are available in Dryad (DOI: 10.5061/dryad.cvdncjtkf). Plasmids generated in this study are available upon request.

## Acknowledgments

We thank Dr. Cassandra Maranas and Sydney Van Gilder for their assistance with experiments; Dr. Maitreya Dunham for feedback on the manuscript; and Dr. Julie Theriot, Dr. Matthew Footer, and Heonsu Kim whose guidance and support were essential for the work with human cells.

This work was supported by the National Institutes of Health (R35-GM148135) and a Washington State Andy Hill Cancer Research Endowment (CARE) Breakthrough Seed Funding Award. JLN is also supported by the Benjamin D. Hall Endowed Chair in Basic Life Sciences.

JOA acknowledges support from the University of Washington Biological Mechanisms of Healthy Aging Training Grant T32AG066574.

