## Supplementary Information for "A conserved dimerization motif squelches transcription"

**Supporting Information for**
### A conserved dimerization motif squelches transcription

Julie M. Ray, Janet Solano Sanchez, Joseph O. Armstrong, Alexander R. Leydon, Jennifer L. Nemhauser

Jennifer L. Nemhauser


**This PDF file includes:**
Figures S1 to S2
Legends for Datasets S1 to S2

**Other supporting materials for this manuscript include the following:**
Datasets S1 to S2

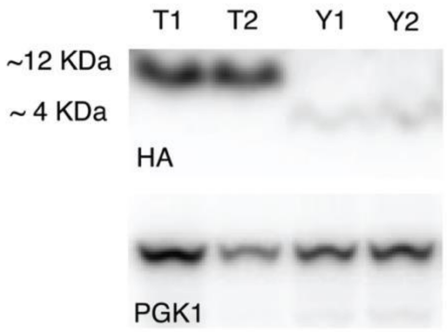


***Fig. S1.*** α-HA western blot of the TPL H1-5 (~12 KDa) and YC5C-H1 (~4 KDa) biological replicates used for RNA-seq, with an α-PGK1 loading control below. All strains are *Saccharomyces cerevisiae.*

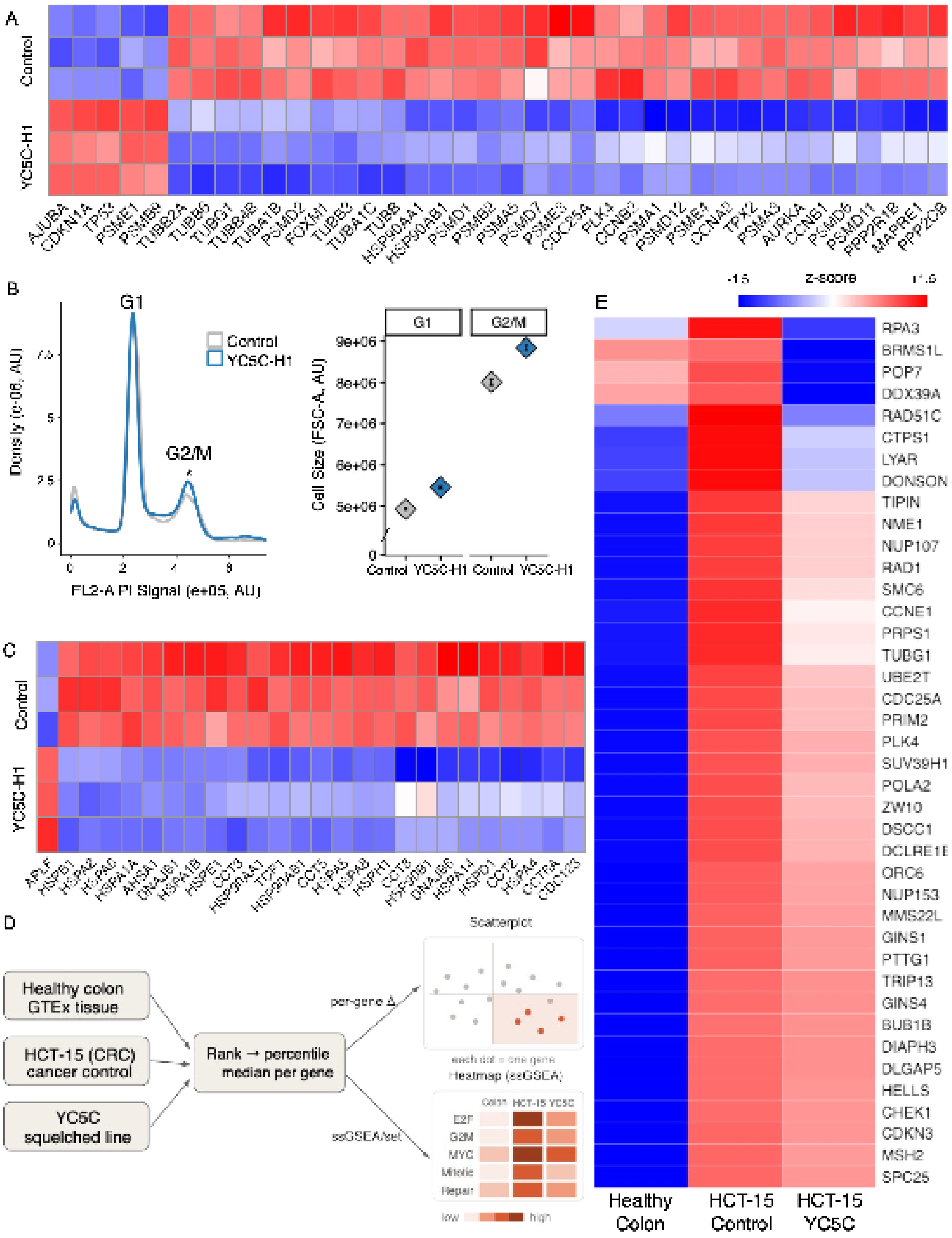


***Fig. S2. YC5C squelches E2F target genes that are highly elevated in CRC. Experiments were performed in the human colorectal cancer line HCT-15; healthy colon expression is from GTEx tissue data.*** (a) Heatmap of G2/M transition genes (padj < 0.001), colored by zscore of expression in three biological replicates each of control and YC5C-H1. Coloring scale: z-score, -1.5 - +1.5. Scales and coloring are identical for parts (a), (c), and (e). (b) Left: PI stain histogram, G1 and G2/M peaks labelled; asterisk denotes significant enrichment in squelched population. Right: Cell sizes (FSC-A, AU) in G1 and G2/M phases. (c) Heatmap of protein folding genes (padj < 0.001). (d) Schematic of the transcriptomic comparison: per-gene percentile ranks from GTEx healthy colon, HCT-15 control and YC5C-squelched HCT-15 are compared gene by gene (scatterplot) and by gene set (ssGSEA heatmap). (e) Hallmark E2F target genes mapped by relative expression in healthy colon tissue, HCT-15 control cells, and squelched HCT-15 cells. Coloring scale: z-score, -1.5 - +1.5. Scales and coloring are identical for G2/M and E2F cancer heatmaps.

**Dataset S1 (separate file).**
Yeast RNA-seq analysis.

**Dataset S2 (separate file).**
Human RNA-seq analysis.

**Supplemental Files on Dryad**
All R scripts for analyzing and plotting data
All unprocessed data (growth assays, cytometry)
All plasmid sequences
